# Transcription factor-promoter interactions for *PSY*/*PSYR*-mediated regulation of root growth

**DOI:** 10.1101/2025.09.29.679155

**Authors:** Jin C.-Y. Liao, Anne-Maarit Bågman, Angie J. Liu, Yejin Shim, Siobhán M. Brady, Pamela C. Ronald

## Abstract

Precise *cis*-regulatory control of gene expression is essential for plant growth. In *Arabidopsis thaliana*, PLANT PEPTIDE CONTAINING SULFATED TYROSINE (PSY) peptides and their receptors (PSYRs) mediate growth-stress trade-offs, yet the transcriptional regulation of these genes remains poorly understood. Here, we mapped transcription factor (TF)-promoter interactions for nine *PSY* and three *PSYR* genes by combining high-throughput enhanced yeast one-hybrid screening with DNA Affinity Purification sequencing (DAP-seq) data, uncovering 1,207 interactions that reveal both shared and gene-specific regulatory relationships, defining the global TF-promoter interaction network of the *PSY*/*PSYR* pathway. Functional analysis of 25 TF mutants identified 12 regulators that significantly influence root growth, most acting as repressors. Of these, CYTOKININ RESPONSE FACTOR 10 (CRF10) emerged as a strong growth inhibitor. We identified a CRF10-binding motif in the *PSYR3* promoter using DAP-seq data and validated it using eY1H. This motif is also located in the last 3′ terminal exon of *Topoisomerase 3A* (*TOP3A*). Guided by these insights, we used CRISPR/Cas9-mediated promoter editing to delete a small region encompassing or flanking a functional TF-binding site (TFBS). Removal of this motif, or of its surrounding region, enhanced root growth, yielding variants that retained root length comparable to the *crf10* mutant. Our results suggest that the observed root growth phenotype results either from disruption of the CRF10 binding motif or from the mutation in the *TOP3A* exon.

## Introduction

Plants continuously face the fundamental challenge of balancing growth and defense, two competing physiological priorities essential for survival and reproduction. Growth maximizes biomass accumulation and competitive ability, while defense mechanisms protect against pathogens and herbivores. This trade-off is tightly regulated by complex signaling networks that integrate environmental cues with internal developmental programs. Small, secreted peptides have emerged as pivotal regulators in this process. The PLANT PEPTIDE CONTAINING SULFATED TYROSINE (PSY) family and their receptors (PSYRs) form a conserved signaling module in *Arabidopsis thaliana* that modulates cell expansion, root development, and various physiological responses^1–5^. PSY peptides promote growth by stimulating cell elongation and division while interfacing with defense pathways to fine-tune resource allocation. Despite their importance, the transcriptional mechanisms controlling *PSYR* and *PSY* gene expression remain poorly defined, limiting insight into how peptide signaling adapts dynamically to environmental and developmental cues. *Cis*-regulatory elements in gene promoters serve as binding platforms for transcription factors (TFs) that orchestrate gene expression spatially and temporally, but integrating high-throughput TF-promoter interaction data into functional regulatory modules remains challenging. Precise manipulation of such *cis*-elements for trait engineering is further complicated by redundancy and incomplete functional annotation.

CRISPR/Cas-mediated promoter and *cis*-regulatory element (CRE) editing provides powerful tools to fine-tune gene expression and optimize crop traits. Approaches include multiplexed promoter targeting, CRE disruption or deletion, and promoter insertion or swapping^6^. Multiplexed CRISPR/Cas tiling screens in tomato, maize, and rice have demonstrated the potential of systematic CRE editing, although large-scale identification of functional elements remains limited^7–9^. Targeted CRE disruption has enhanced stress responses and disease resistance. In rice, editing the GT-1 element (‘GAAAAA’) in the *Oryza sativa RAV2* (*OsRAV2*) promoter modulated salt tolerance^10^, while deletion of a TALe (Transcription Activator-Like effector)-binding element in the *SWEET11*, a sugar transporter gene, promoter improved bacterial blight resistance without affecting fertility^11–13^. Similarly, mutations in barley’s *Hordeum vulgare* purple acid phosphatase a (*HvPAPhy_a*) promoter reduced phytase activity^14^, and disruption of Lateral Organ Boundaries 1 (LOB1) promoter elements in citrus conferred canker resistance^15–17^. CRE editing in introns or downstream regions further expands versatility, as shown by gain-of-function *Solanum lycopersicum* WUSCHEL (*SlWUS*) alleles in tomato^7,18^. Beyond plants, CRISPR-mediated disruption of transcription factor binding sites has been demonstrated in human cells and preclinical models^19^. In crops, understanding transcriptional networks that regulate growth-defense trade-offs, where activating defense often limits growth, will be critical for engineering resilient plants with optimized productivity.

Here, we integrated high-throughput enhanced yeast one-hybrid (eY1H) screening of nine *PSY* and three *PSYR* promoters with DAP-seq data to generate a comprehensive TF-promoter interaction network comprising over 1,200 interactions. Functional validation of selected TFs through loss-of-function mutant alleles and overexpression lines identified regulators modulating *PSY/PSYR* expression and root system architecture (RSA) phenotypes. Informed by this network, we used CRISPR/Cas9-based promoter editing to mutate a CRF10-binding motif located within the last exon of *Topoisomerase 3A*, which overlaps the *PSYR3* promoter region. These mutations led to measurable changes in *PSYR3* expression. Together, our findings establish a framework for mapping and manipulating the transcriptional regulation of the *PSY*/*PSYR* signaling pathways.

## Results

### Mapping the putative *PSY/PSYR* transcriptional regulatory network

To investigate transcriptional regulation underlying PSY peptide signaling, we performed enhanced yeast one-hybrid (eY1H) assays on three *PSYR* and nine *PSY* promoters using a curated *Arabidopsis* transcription factor (TF) consisting of 2,000 TFs^20^. Publicly available DAP-seq datasets^21^ describe 1,207 TF-promoter interactions with ∼500 TFs assessed. When integrated, these interactions revealed both shared and gene-specific regulatory relationships across the *PSY/PSYR* family, forming a comprehensive predictive transcriptional map controlling PSY and peptide transcription (Fig. 1 and Dataset S1). Network analysis further uncovered clusters of TFs potentially coordinating growth and defense responses, and several central hubs emerged as candidate regulators for functional validation. eY1H assays identify promoter-TF interactions in a chromatinized context in yeast, which can reveal potential regulatory relationships but may include false positives and negatives. Published studies have shown that a subset of eY1H-identified interactions is supported by *in planta* validation, such as ChIP or genetic analysis^22,23^. In contrast, DAP-seq profiles interactions between individual TFs and DNA sequences *in vitro*, providing high-resolution information on potential binding sites but not direct evidence of *in vivo* occupancy^21^. Integration of both datasets revealed limited overlap between the two datasets. Specifically, three TFs were identified to interact with the *PSYR1* promoter, eight for *PSYR2*, six for *PSYR3*, sixteen for *PSY1*, four for *PSY2*, two for *PSY3*, seven for *PSY4*, one for *PSY5-8*, and three for *PSY9* (Fig. S1 and Dataset S2). Together, these shared interactions highlight a small but robust set of candidate regulators supported by both approaches^24^.

**Fig. 1.**
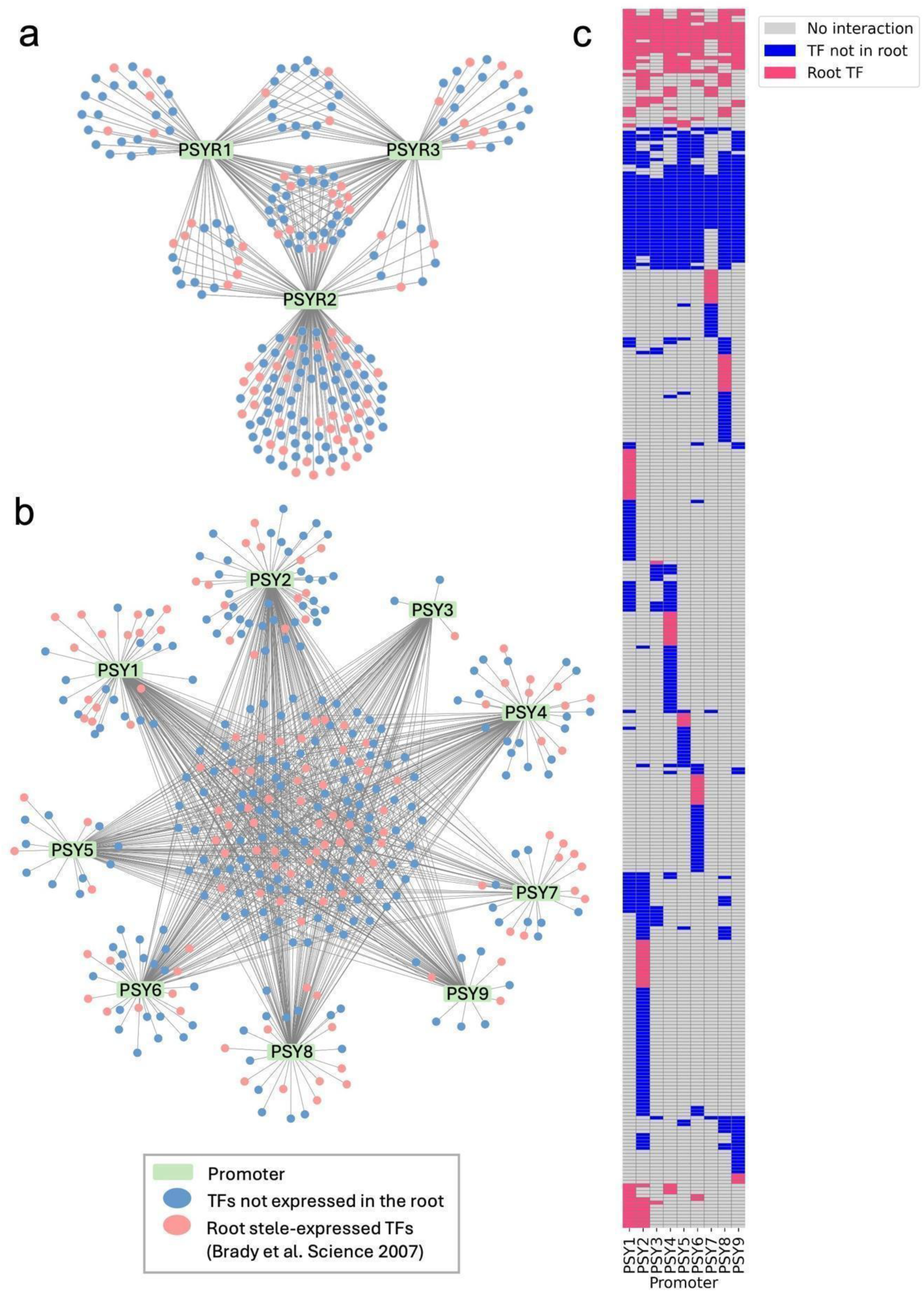
TF-*PSY/PSYR* promoter interaction network. **a.** TF-promoter interactions involved in PSY perception, identified by eY1H (*PSYR1-PSYR3*, 3 promoters). **b.** TF-promoter interactions involved in PSY peptide transcription, identified by eY1H (*PSY1-PSY9*, 9 promoters). Rectangles represent promoters; small circles represent TFs. Pink circles indicate root-expressed TFs (Brady et al., 2007), and blue circles indicate TFs outside the root stele-expressed set. **c.** Heatmap of TF-promoter interactions for PSY peptide transcription identified by eY1H, with shared TFs clustered. Coloring follows panel b: pink = root-expressed TFs (Brady et al., 2007), blue = TFs outside the root stele-expressed set.

To investigate how these regulatory interactions are reflected *in planta*, we examined gene expression patterns across tissues and cell types. Analyses of publicly available RNA-seq datasets^25^ defined the expression profiles of *PSYR* and *PSY* genes throughout Arabidopsis organs. All three *PSYR*s were broadly expressed, with *PSYR1* enriched in shoots and *PSYR3* in roots (Fig. S2). Among the peptides, *PSY1* was the most abundant and widely expressed, whereas *PSY7* showed minimal expression. *PSY4* was predominantly localized to roots, and *PSY9* was mainly expressed in seeds (Fig. S2). Single-cell data from the Plant sc-Atlas^26^ further revealed cell-type specificity. *PSY3* and *PSY4* co-localized in root cell types, suggesting overlapping functions, while *PSY6* and *PSY8* displayed distinct but complementary root-specific patterns (Fig. S3). Together, these findings highlight potential links between spatial regulation, receptor-ligand interplay, and root system architecture (RSA).

### Network-guided selection and screening of TFs for promoter editing

To systematically prioritize TFs for functional analysis, we combined whole-promoter interaction assays (eY1H) with DAP-seq data, identifying 40 high-confidence candidates. From these, 12 TFs with available homozygous mutant lines and an additional 10 TFs detected only by DAP-seq (represented by 13 independent homozygous alleles) were selected for phenotypic analysis (Dataset S2, S3, Table S2). Using these prioritized TFs, we then mapped their respective transcription factor binding sites (TFBS) across all three *PSYR* and nine *PSY* promoters to identify precise promoter regions for targeted editing. We focused on root system architecture (RSA) as a readout, given its responsiveness to genetic and environmental cues and the availability of high-throughput phenotyping tools. We then developed a workflow integrating TFBS mapping with functional assays to pinpoint regulatory regions suitable for CRISPR-based promoter editing (Fig. 2a). Although a root-specific DAP-seq dataset would be ideal, it is not currently available, underscoring the importance of combining multiple data sources for candidate prioritization.

**Fig. 2.**
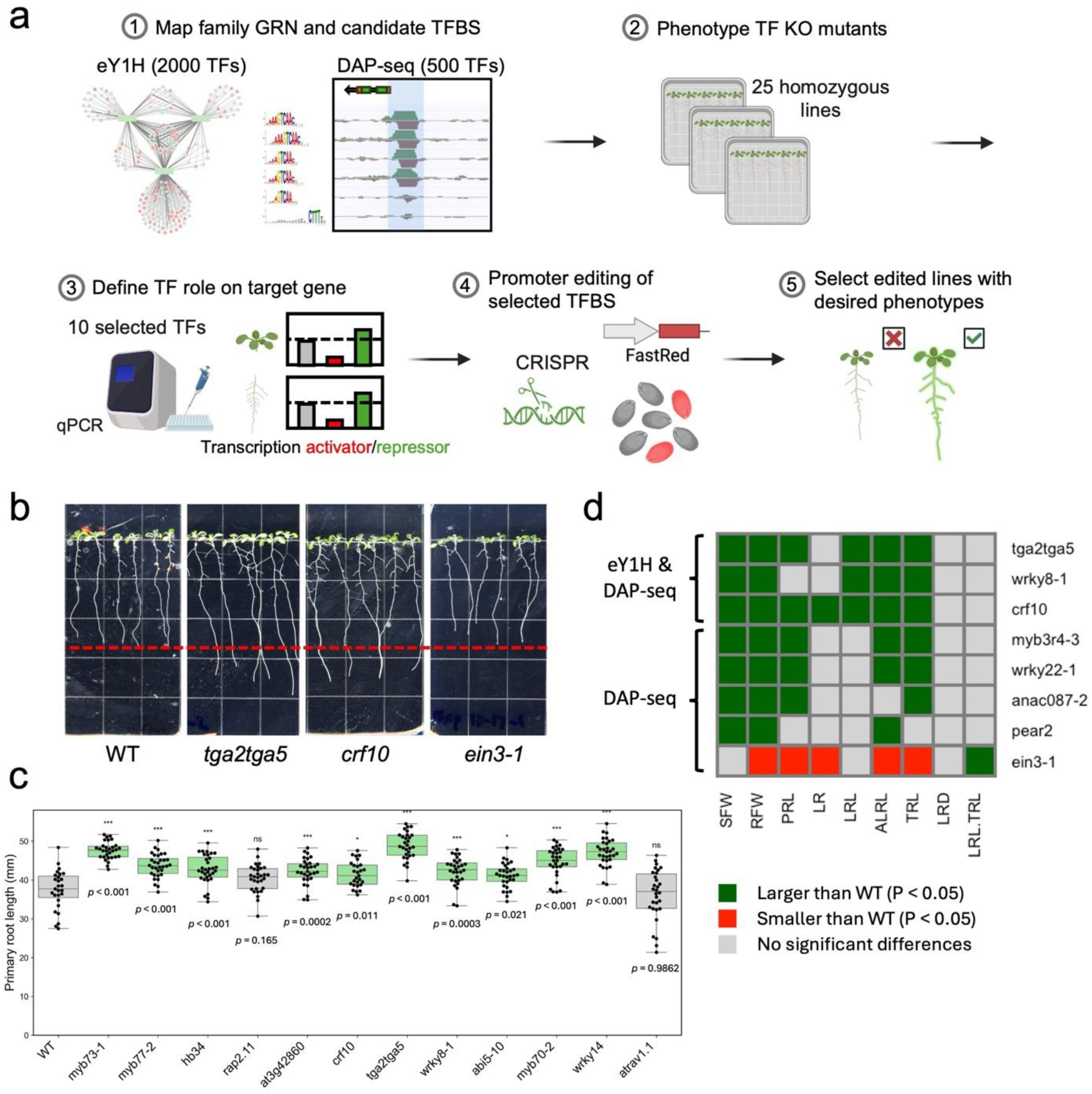
Identification of putative TFBS. **a.** Schematic of the screening and editing workflow applied to *PSY/PSYR* promoters. **b.** Representative 10-day-old TF knockout seedlings identified based on validated TF-promoter interactions. WT, Col-0. **c.** Primary root length of 12 TF knockout mutants at 10 days old. Primary root length of 12 TF knockout mutants at 10 days old. At least 20 plants per genotype were analyzed in each experiment, and the experiment was independently repeated three times. Green indicates a statistically significant increase relative to WT, and grey indicates not significantly different (*** *P* < 0.001, ** *P* < 0.01, * *P* < 0.05, one-way ANOVA; exact n is in Supplementary Data 3). **d.** Heatmap showing phenotypes associated with TF knockout alleles. Selected mutant alleles, identified from both eY1H and DAP-seq or from DAP-seq only, are listed in rows, and measured traits are shown in columns. Statistically significant differences relative to WT are indicated by colored cells (*P* < 0.05, one-way ANOVA; exact n and *P* values are in Supplementary Data 3). From left to right, traits include shoot fresh weight (SFW, g per 10 seedlings), root fresh weight (RFW, g per 10 seedlings), primary root length (PRL, mm), number of lateral roots (LR), total lateral root length (LRL, mm), total root length (TRL, PRL + LRL, mm), average lateral root length (ALRL, LRL/LR, mm), lateral root density (LRD, LR/PRL), and proportion of total root length contributed by lateral roots (LRL/TRL). Root traits were measured from 10-day-old seedlings. Green indicates a statistically significant increase relative to WT, red indicates a statistically significant decrease, and grey indicates not significantly different from WT (*P* < 0.05, one-way ANOVA; exact n and *P* values are in Supplementary Data 3).

While *PSY1* overexpression and the *psyr1,2,3* triple mutant have been reported to alter primary root length^1–3,27^, other RSA phenotypes in these lines, as well as in single *PSY* or *PSYR* mutants, remain unknown. Given the root cell type expression patterns of *PSY/PSYR* genes, we focused on phenotypic analysis of mutants for the 12 TFs identified by both eY1H and DAP-seq. Knockouts were first validated by PCR genotyping and qRT-PCR (Fig. S5). Single mutant alleles of ten TF mutants (*crf10*, *tga2 tga5*, *wrky8-1*, *myb77-2*, *myb73-1*, *hb34*, *myb70-2*, *at3g42860*, *abi5-10*, *wrky14*) displayed increased primary root length, while two mutants (*rap2.11*, *atrav1.1*) showed no change (Fig. 2b-d). In addition to these 12 TFs, we also analyzed DAP-seq-only candidates for phenotypic effects, as homozygous mutant lines were available and they could reveal additional regulators of root growth not detected by eY1H. Among the DAP-seq-only TFs, six alleles of four TFs (*wrky22-1, wrky22-2, wrky22-3, anac087-2, erf11, aif*) displayed increased root length, one allele (*ein3-1*) showed decreased root length, and seven alleles (*hca2, pear2, myb3r4-3, tga10, ntl4-1, erf4-1, ntl8-1*) exhibited no change (Fig. S6). Many of these alleles had been studied previously, but root phenotypes were largely unreported or poorly described, with *myb70-2* as the only clearly documented case^28^, showing a longer primary root consistent with our observations. Overall, 26 mutants of 24 TFs exhibited significant changes in root growth compared with wild-type plants, 16 of which displayed increased biomass and enhanced root growth, suggesting that multiple TFs negatively regulate growth to fine-tune developmental outcomes (Data S3). We tested the less-characterized candidate CRF10 and found that overexpression caused stunted growth, whereas the *crf10* knockout displayed enhanced growth compared with wild type (Fig. 2 b-d and Fig. S7). Collectively, these results suggest that the identified TFs contribute to root growth regulation, highlighting the potential of promoter-targeted approaches to modulate RSA phenotypes.

### Transcript levels in shoots and roots of selected TF knockout lines

To assess whether candidate TFs act as activators or repressors of *PSY/PSYR* transcription and whether their regulation is organ-specific, we measured transcript levels in shoots and roots of selected TF knockout lines using quantitative RT-PCR (qRT-PCR) (Fig. 3). The same homozygous knockout lines that were genotyped and phenotypically validated in Fig. 2c were used for these expression assays, ensuring that transcriptional differences could be directly linked to previously observed RSA phenotypes. We focused on the three *PSYR* and three *PSY* (*PSY1*, *PSY2*, *PSY4*) promoters with the highest number of candidate TF interactions identified by eY1H, and for which binding sites were also detected in DAP-seq (Fig. 3). Shoot and root organs from these validated lines were collected separately to capture potential organ-specific regulation. qRT-PCR analyses revealed consistent significant differences in *PSYR* and *PSY* expression (Fig. 3), directly connecting TF-promoter interactions to transcriptional outputs.

**Fig. 3.**
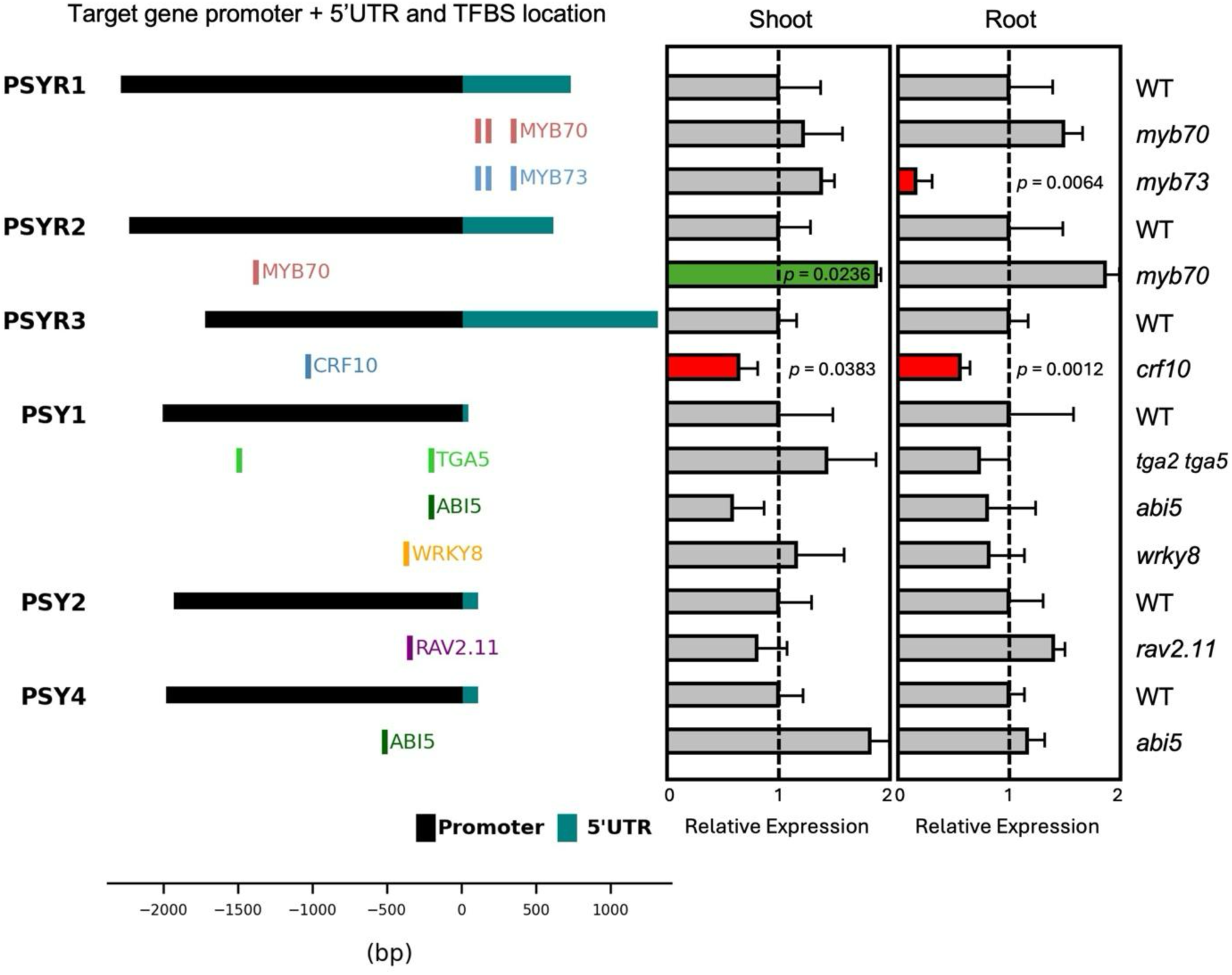
Organ-specific transcriptional changes in TF knockout mutants. Expression of targeted *PSY/PSYR* genes in seedling shoots and roots of 10-day-old TF knockout seedlings. Left, schematic of target *PSY/PSYR* promoters and 5’UTR showing the locations of transcription factor binding sites (TFBS) validated via DAP-seq. Right, relative expression of targeted *PSY/PSYR* genes in TF knockout seedlings. For each biological replicate, 10 shoots or 20 roots were pooled. Gene expression was measured by qRT-PCR, with WT normalized to 1 (grey bars). Expression levels significantly increased or decreased in the TF knockout relative to WT are indicated in green (TF is a transcriptional repressor) or red (TF is a transcriptional activator), respectively. Values represent mean ± SD (n = 3 biological replicates).

Among the analyzed mutants, *myb73*, *myb70*, and *crf10* showed significantly altered expression of *PSY/PSYR* target genes, with distinct patterns of activation and repression. MYB70 bound the promoters of both *PSYR1* and *PSYR2* and repressed *PSYR2* transcription, as loss of MYB70 led to a ∼1.9-fold increase relative to WT (*p* = 0.0236) in the shoot, whereas MYB73 bound the *PSYR1* promoter and activated *PSYR1* transcription, as loss of MYB73 reduced expression to ∼0.14-fold relative to WT (*p* = 0.0064) in the root. CRF10 bound the *PSYR3* promoter and activated transcription in both organs, as loss of CRF10 reduced expression to ∼0.64-fold relative to WT in the shoot (*p* = 0.0383) and ∼0.53-fold relative to WT in the root (p = 0.0012). Several mutants of additional TFs, including TGA5, WRKY8, ABI5, and RAV2.11, were tested for effects on *PSY*/*PSYR* gene expression (Fig. 3 and Data S4), but their precise regulatory roles remain unclear and will require further analysis, including more biological replicates and other mutant alleles. The transcriptional changes observed in *myb70*, *myb73*, and *crf10* are consistent with direct TF-mediated regulation, and the magnitude and organ specificity of these changes corresponded with the root phenotypes of the respective knockout lines, suggesting that these TFs may contribute to fine-tuning *PSY*/*PSYR* signaling.

### Mutation of the CRF10 binding motif in the *PSYR3* promoter/*TOP3A* final exon affects root growth in genome-edited lines

To test whether disruption of transcription factor binding sites could modulate gene expression and plant growth, we implemented a CRISPR/Cas9-based promoter editing strategy. Candidate motifs within *the PSYR3* promoter were targeted, which overlap with the last exon of the *Topoisomerase 3A* (*TOP3A*) coding region. A putative CRF10 binding motif (‘CGGCGG’) was identified in the *PSYR3* promoter using DAP-seq data and validated by eY1H. CRF10 is a member of the APETALA2 (AP2)/ETHYLENE RESPONSE FACTOR (ERF) transcription factor family, which regulates hormone signaling, stress responses, and development^29^. Although the *Arabidopsis* CRF family contains 12 members, CRF10 remains largely uncharacterized. In Col-0 plants, *CRF10* was expressed ubiquitously, with the highest expression in embryos (Fig. S4). In our screening (Fig. 2), *crf10* mutants exhibited longer primary roots and overall enhanced root system architecture (RSA), characterized by increased lateral root number and length as well as greater total root length (Fig. 2b-d). Overexpression of *CRF10* in wild-type Col-0, in two independent lines, caused growth inhibition, confirming that *CRF10* functions as a negative regulator of growth (Fig. S7).

To test whether CRF10 binding influences *PSYR3* expression, we generated CRISPR-edited lines targeting the CRF10 motif in the *PSYR3* promoter/*TOP3A* final exon. Two guide RNAs were designed to target both the primary TFBS and adjacent sequences (Fig. 4a). Sequencing of independent lines revealed two distinct alleles. R3pro_Var1 carries a 31 bp deletion immediately adjacent to the DAP-seq-validated CRF10 binding motif, leaving the consensus sequence intact, whereas R3pro_Var2 carries a 27 bp deletion that removes the motif entirely. T2 homozygous lines were used for phenotyping and expression assays. Notably, R3pro_Var1 and R3pro_Var2 alleles are also present in the last 3′ terminal exon of *Topoisomerase 3A* (*TOP3A*), raising the possibility that the phenotypes observed in the promoter-edited variants may be due to the TOP3A mutation (Fig. 4a). The *top3a-1* mutant carries a T-DNA insertion that abolishes mRNA spanning the insertion site, resulting in a knockout allele with severe developmental defects or early lethality^31^. *top3a-2* carries a T-DNA insertion upstream in the gene, producing a hypomorphic allele that is viable but completely sterile, with milder growth abnormalities such as dwarfing and organ curling^30,31^. In the absence of available phenotypes for mutations in the last exon of *TOP3A*, it is unknown whether such mutations confer the same phenotype observed in our study.

**Fig. 4.**
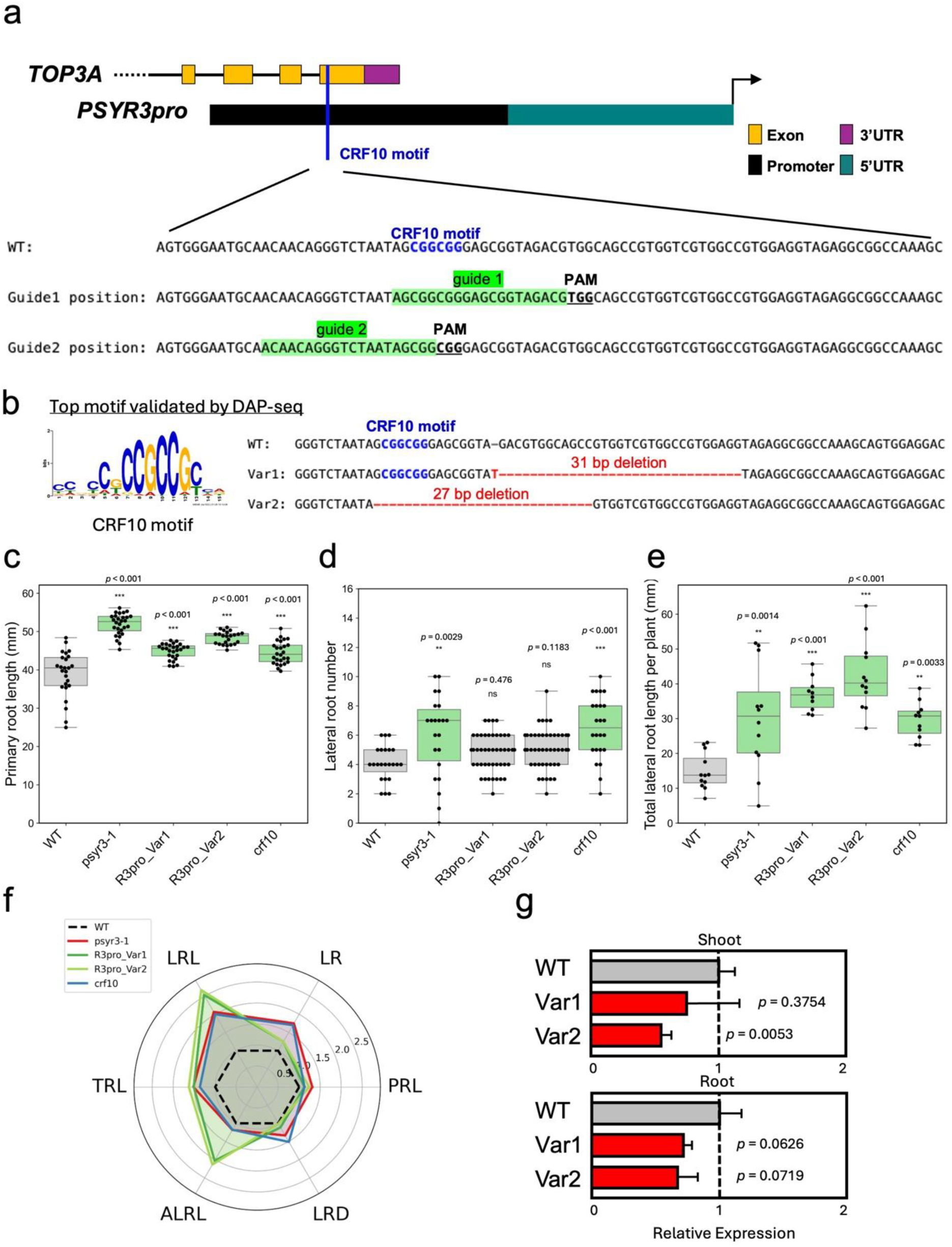
CRF10 motif editing in the *PSYR3* promoter and the *TOP3A* last exon. **a.** Schematic of the *PSYR3* promoter and *TOP3A* 3’ exons showing the CRF10 motif targeted for deletion. **b.** DNA sequence comparison of two promoter-edited variants (Var1 and Var2) relative to WT. Deleted CRF10 motifs are highlighted in red; dashes indicate deleted nucleotides. **c-f.** Root traits measured in 10-day-old seedlings. At least 20 plants per genotype were analyzed in each experiment, and the experiment was independently repeated three times. Green indicates a statistically significant increase relative to WT, and grey indicates not significantly different (*** *P* < 0.001, ** *P* < 0.01, * *P* < 0.05, one-way ANOVA; exact n is in Supplementary Data 3). **c.** Primary root length of WT, *psyr3-1*, *crf10*, and promoter-edited plants. Boxplots show median, interquartile range, and full range. **d.** Number of visible lateral roots per genotype. **e.** Total lateral root length per genotype. **f.** Radar plot comparing root traits across genotypes: PRL, primary root length (mm); LR, number of lateral roots; LRL, total lateral root length (mm); ALRL, average lateral root length (LRL/LR, mm); TRL, total root length (PRL + LRL, mm); LRD, lateral root density (LR/PRL). **g.** *PSYR3* expression in shoot and root organs of 10-day-old promoter-edited seedlings. Values represent mean ± SD (n = 3 biological replicates).

Phenotypic analysis revealed that promoter-edited plants exhibited altered primary root length compared with wild type, with phenotypes comparable in magnitude to *psyr3-1* and *crf10* knockout mutants (Fig. 4c). Lateral root number remained unchanged (Fig. 4d), but total lateral root length per plant increased (Fig. 4e), resulting in increased total and average lateral root length relative to WT, *psyr3-1*, and *crf10* (Fig. 4f). Both edited alleles showed a mild reduction in *PSYR3* transcript abundance in shoots and roots, with R3pro_Var1 and R3pro_Var2 exhibiting ∼0.78- and ∼0.56-fold expression relative to wild-type controls (Fig. 4g). Although these decreases were not statistically significant in R3pro_Var1 (shoot: *P* = 0.3754; root: *P* = 0.0626) or in the root of R3pro_Var2 (*P* = 0.0719), the shoot of R3pro_Var2 showed a significant reduction (*P* = 0.0053), with the CRF10 consensus motif removed. This trend suggests that CRF10 consensus motif and sequences adjacent to the motif may contribute to transcriptional regulation, although effects from the linked *TOP3A* mutation cannot be excluded.

## Discussion

Our work demonstrates the utility of network-guided promoter analysis to identify putative TF binding sites. We observed altered root growth phenotypes in plants where the CRF10 binding motif itself was deleted, and when nearby promoter regions were perturbed. Notably, the CRF10 binding motif we analyzed is also located in the last 3′ terminal exon of *Topoisomerase 3A* (*TOP3A*). Thus, we cannot rule out the possibility that the observed root growth phenotypes are also due to the mutated TOP3A.

TF-promoter interaction analysis of the *PSY*-*PSYR* signaling pathway provides a framework for future precise, context-aware trait modulation. By leveraging functional *cis* elements identified from regulatory networks, edits can be tailored for spatially precise regulation at the organ, tissue, or even cell level, in addition to being environment-responsive, thereby enhancing trait specificity while minimizing unintended systemic effects. This capability is particularly valuable for complex traits influenced by multiple environmental and developmental cues, where traditional knockout approaches may produce broad pleiotropic effects. Future work will be directed at genome editing of experimentally validated TF binding site motifs with the aim of optimizing traits while preserving overall plant fitness.

Although tiling-deletion-based screens have enabled systematic functional interrogation of regulatory regions, they are labor-intensive, time-consuming, and often less predictive, particularly when applied across multiple genes. Moreover, because these strategies rely on scanning promoter regions through serial deletion or mutagenesis, they tend to reveal functional elements retrospectively rather than enabling targeted modulation. In contrast, our gene regulatory network-guided strategy enables identification of putative functional *cis*-regulatory elements. This hypothesis-driven approach sets up a framework for future fine-tuning of gene expression. By narrowing the range of potential target sites, network-guided predictions allow the researcher to design genome edits that can enhance both predictability and efficiency. Although promoter insertions or swaps remain technically challenging, successful examples, such as the insertion of the *GOS2* promoter upstream of *ARGOS8* in maize to enhance expression and improve drought tolerance^32^, demonstrate the potential for context-aware regulatory engineering and create opportunities for synthetic promoter applications by replacing native promoters.

Promoter editing in Arabidopsis remains largely unexplored, yet the species offers rich genomic and regulatory resources, including well-characterized gene regulatory networks, TFBS maps, and promoter motifs. Comparative analyses between Arabidopsis and crop species will be essential to translate these insights to agricultural contexts. While gene regulatory networks are extensively mapped in Arabidopsis, coverage in major crops remains limited. Systematic comparisons of TFBSs, promoter architectures, and regulatory modules can identify conserved elements, enabling rational design of network-guided promoter edits in Arabidopsis and crop species. Future studies will explore whether promoter edits of putative TFBSs in the *PSY*/*PSYR* promoters affect RSA and pathogen resistance. Integrating ATAC-seq or other chromatin profiling approaches in edited lines will provide deeper insight into the epigenomic consequences of TFBS-level perturbations and help refine predictive models of *cis*-regulatory engineering.

## Methods

### Promoter cloning for eY1H

*PSYR* and *PSY* gene promoters previously tested in Ogawa-Ohnishi et al.^1^ were cloned using the method described by Gaudinier et al.^22,33^. Promoters were amplified by PCR from *Arabidopsis thaliana* Col-0 genomic DNA using Phusion High-Fidelity DNA Polymerase (NEB). The amplified promoter fragments were first cloned into the pENTR 5’TOPO vector using the pENTR 5’TOPO kit (Invitrogen) and fully sequenced. Subsequently, promoters were recombined with the Gateway LR reaction into both pMW2 and pMW3^41^ Gateway destination vectors (designed for yeast expression and containing respectively *HIS3* or *LacZ* reporter genes) using LR Clonase™ II (Invitrogen). The resulting constructs were sent to Azenta Life Sciences (South Plainfield, NJ, USA; formerly GeneWiz) for whole plasmid sequencing using the Plasmid-EZ nanopore-based sequencing method.

### Enhanced Yeast One Hybrid (eY1H) Assay

Assays were performed according to Gaudinier et al.^22,33^. *PSYR* and *PSY* promoter – pMW2/pMW3 reporter constructs were transformed into yeast strain YM4271. Resulting yeast colonies were screened for autoactivation for both histidine biosynthesis and lacZ activity, and the construct presence within the yeast genome was confirmed by PCR-based genotyping. The resulting bait yeast strains were screened against a complete collection of 2000 Arabidopsis transcription factors^34^ in a mating-based assay, and positive interactions were identified for both histidine and lacZ activity compared to a negative control pDESTAD-2μ. An interaction was recorded as successful if there was activation of either HIS or lacZ. The eY1H Yeast One-Hybrid screening was carried out by the Yeast One-Hybrid Services Core at the UC Davis Genome Center, at the University of California, Davis. The table with all the results can be found at https://github.com/jcyliao/Liao2025.

### Network construction

Networks were made using Cytoscape v.3.10.2^35^. All cytoscape network files can be found at https://github.com/jcyliao/Liao2025.

### DAP-seq data comparison and mutant lines selection

DAP-seq TF binding data generated by O’Malley et al.^21^ were used to identify direct TF-promoter interactions. DAP-seq target genes (fraction of reads in peaks [FRiP] ≥ 5%) for 529 TFs were downloaded from the Plant Cistrome Database. TFs binding to each individual *PSY/PSYR* promoter were extracted from the DAP-seq data (Dataset S1). A Python script was then used to compare the TF lists for each promoter from eY1H and DAP-seq experiments, generating lists of overlapping TFs identified by both assays (Dataset S2).

From these comparisons, 40 overlapping TFs and ∼20 TFs identified only in DAP-seq were selected based on the availability of loss-of-function mutants. Mutant lines previously used in published studies were prioritized (Table S2).

### Expression analysis of *PSY/PSYR* genes and their regulatory TFs

Expression patterns of *PSYR* and *PSY* genes, along with their selected potential regulatory transcription factors (TFs) identified from eY1H and DAP-seq, were analyzed using publicly available datasets. Bulk RNA-seq datasets were collected to examine gene expression across different tissues and developmental stages. Single-cell RNA-seq datasets, including root cell type-specific profiles from the Plant sc-Atlas database^26^, were used to assess cell type-specific expression. Comparative analyses were performed to identify co-expression patterns and potential tissue- or cell type-specific regulatory relationships.

### Plant materials and growth conditions

All transgenic lines were generated in the *Arabidopsis thaliana* Col-0 accession, except for *abi5-1*, which was in the Ws-2 background. Loss-of-function TF mutants were obtained from the Arabidopsis Biological Resource Center (ABRC) at Ohio State University. All mutant lines, including those carrying T-DNA insertions, fast neutron-induced mutations, or EMS-induced mutations, were genotyped to confirm homozygosity. Quantitative RT-PCR (qRT-PCR) was performed to verify transcript-level alterations (Fig. S5 and Data S4). The primers used for genotyping and qRT-PCR, along with accession numbers and additional details on the analyzed mutant lines, are provided in Supplementary Table 2. Plants were grown under a 16-hour light/8-hour dark photoperiod for propagation.

### Generation of TF overexpressing transgenic plants

The coding sequences for each TF used in this study were obtained from the TF ENTR clone collection^34^ and subsequently cloned into the N-GFP tag pGWB506 (Addgene #74848) destination vector using LR Clonase™ II (Invitrogen) for overexpression in plants. The resulting GFP-TF constructs, driven by the 35S promoter, were introduced into *Agrobacterium tumefaciens* strain GV3101 and used to transform *A. thaliana* (Col-0) via the floral dip method^36^. At least ten independent transgenic lines were selected on ½ MS solid medium supplemented with 25 mg/L hygromycin. Transgene expression was assessed through GFP fluorescence microscopy and PCR. For each genotype, two to three independent T3 homozygous plants were used for subsequent phenotypic analyses. The primers used for genotyping and accession numbers, and additional details on the constructs, are provided in Supplementary Table 1.

### Plant phenotype analysis of the TF mutants

Homozygous TF loss-of-function mutants, TF overexpression T3 plants, and the wild-type (WT) controls were sterilized with 25% (v/v) bleach for 20 min, followed by 5 washes with sterile Milli-Q water. Sterilized seeds were plated on ½ MS medium supplemented with 1% sucrose and 0.6% Gelzan, then stratified at 4°C for 2-3 days before being transferred to a growth chamber under long-day (LD) conditions (16 h light/8 h dark) with 50-60% humidity at 22°C. Mutant lines and wild-type controls were arranged in a randomized block design within the growth chamber. Plate images were scanned at 10 days post-germination (10 dpg) using an Epson Expression 12000XL scanner at 300 dpi resolution. Root traits were measured from the scanned images, with at least 20 plants per genotype included in each experimental replicate. The experiment was repeated independently three times. For root trait measurements, RootNav 1.0^37^ was used for image analysis and data extraction. Traits measured included primary root length (PRL), lateral root number (LR) and lateral root length (LRL). Composite traits were analyzed, including total root length (TRL = PRL + LRL), average lateral root length (ALRL = LRL/LR), lateral root density (LRD = LR/PRL), and the percentage of LRL contributing to TRL (LRL/TRL). Variation across these mutants relative to WT was assessed using one-way ANOVA for all RSA traits. Statistical analyses were performed in Python, with significance determined by Tukey’s HSD test.

### RNA extraction and quantitative PCR

The whole root and shoot parts of 10-day-old seedlings were collected separately. For one shoot sample, shoots from 10 seedlings grown on the same plate were pooled together. For one root sample, roots from 30 seedlings grown on the same experiment were pooled together. Three independent replicates per genotype were collected. Total RNA was extracted using TRIzol reagent (Invitrogen) for TF mutant screening experiments and using the RNeasy Plant Mini Kit (QIAGEN) for promoter-edited line experiments, following the manufacturer’s instructions. Genomic DNA contamination was removed by RNase-free DNase treatment during RNA extraction, according to the manufacturer’s protocols. First-strand cDNA was synthesized using the SuperScript™ III First-Strand Synthesis System for RT-PCR (Invitrogen) following the manufacturer’s instructions. Quantitative real-time PCR (qRT-PCR) was performed using iTaq™ Universal SYBR Green Supermix (Bio-Rad) on a Bio-Rad CFX96 Real-Time System coupled with a C1000 Thermal Cycler (Bio-Rad) under the following conditions: initial denaturation at 95°C for 30 s, followed by 39 cycles of PCR (denaturation at 95°C for 5 s, annealing at 60°C for 30 s, and extension at 72°C for 20 s). Relative gene expression was determined using the 2^−ΔΔCT^ method, with ACTIN2 as the endogenous control. The primers used for qRT-PCR are provided in Supplementary Table 1.

### Generation of CRISPR/Cas9 promoter editing of CRF10 binding site and plant screening

Promoter regions containing characterized TFBSs were selected based on TF-promoter interaction mapping and mutant phenotypes. Guide RNAs (gRNAs) were designed to flank these sites, and corresponding CRISPR/Cas9 constructs were generated using standard cloning protocols.

For the generation of pJL-AtU6.26-gPSYR3-S1S2 entry clone, an EcoRI-AtU6-26 promoter-partial pQZ435–BsaI gBlock was synthesized via the IDT gBlock service. gRNAs were designed using CHOPCHOP^38^, and PTG assembly into pQZ435, with the OsU3 promoter removed, was performed following Xie et al.^39^ with minor modifications. The entry clone was transferred into the pRU294 destination vector via Gateway LR recombination, following the manufacturer’s instructions, and the resulting construct was transformed into *Agrobacterium tumefaciens* GV3101 for plant transformation. Constructs were transformed into *Arabidopsis thaliana* via Agrobacterium-mediated floral dipping. Rapid screening of Cas9-containing and Cas9-free lines in both T1 and T2 generations was conducted using a seed coat fluorescence marker^40^. Transgenic plants were confirmed by PCR and Sanger sequencing to confirm precise edits disrupting TFBSs. Successfully edited lines were propagated for subsequent phenotypic and gene expression analyses.

### Figure preparation and statistical analysis

Python was used for statistical analyses and to generate plots. Figures were assembled in Microsoft PowerPoint. Some illustrations in the workflow (Fig. 2a) were created using BioRender.

## Data availability

All supplementary datasets, Cytoscape files, and Python code are available at https://github.com/jcyliao/Liao2025.

## Use of AI

ChatGPT (OpenAI) assisted only with improving the phrasing of some paragraphs and did not generate any new content.

## Acknowledgements

We thank Bhanu Upadhyayula, Kai Deshpande, and Maya Ziebarth for assistance with organizing genetic materials and performing genotyping experiments. We are also grateful to Valley Stewart for guidance and mentoring, and to Qiong Zhang for providing the pQZ435 vector and the modified PTG cloning protocol. This work was supported by the Chan Zuckerberg Initiative and NIH MIRA grant #R35 GM148173 to P.C.R. We thank Idan Efroni for critical reading of the manuscript and for helpful suggestions.

## Author contributions

J.C.-Y.L. conceived the project, designed the experiments, and conducted most of the experiments and all data analysis. A.B. and the Yeast One-Hybrid (Y1H) Services Core conducted all eY1H screenings. J.C.-Y.L. and A.J.L. prepared samples for the gene expression assays. A.J.L. performed most of the root phenotyping raw data acquisition and, together with Y.S., carried out qRT-PCR measurements. Both A.J.L. and Y.S. participated in mutant screening. P.C.R. supervised the work and provided funding and resources. J.C.-Y.L. wrote the manuscript. P.C.R. and J.C.-Y.L. revised the MS with input from S.M.B.

## Supplemental Materials

All supplemental materials referenced in the text will be provided in the final published version. All main findings are fully presented in the figures included in this manuscript.

## Supplementary Figures

**Supplementary Fig. 1. Overlap of eY1H and DAP-seq interactions.**

Venn diagram showing TFs identified by either eY1H or DAP-seq for three *PSYR* promoters and nine *PSY* promoters. Yellow circle indicates TFs identified by eY1H, and blue circle indicates TFs identified by DAP-seq.

**Supplementary Fig. 2. *PSY/PSYR* expression at the whole plant level.**

Data were extracted from the publicly available Arabidopsis RNA-seq (ARS) database (http://ipf.sustech.edu.cn/pub/athrna/).

**Supplementary Fig. 3. *PSY/PSYR* expression at the single-cell level in roots.**

Data were extracted from the Plant sc-Atlas database.

**Supplementary Fig. 4. Expression patterns of candidate TFs identified by both eY1H and DAP-seq.**

Data were extracted from the publicly available bulk RNA-seq database.

**Supplementary Fig. 5. Confirmation of selected TF knockouts.**

qRT-PCR analysis of selected TF expression in TF knockout (KO) mutants. Gene expression was measured by qRT-PCR and normalized to wild-type (WT = 1, grey bars). Expression levels showing a significant decrease are indicated in red. Values represent mean ± SD (n = 3 biological replicates).

**Supplementary Fig. 6. Comparison of primary root length in selected Arabidopsis mutant alleles of transcription factors identified by DAP-seq.**

Primary root length of 10 TF KO mutants at 10 days old. Exact *P* values are reported in Supplementary Data 3.

**Supplementary Fig. 7. CRF10 acts as a negative regulator of plant growth.**

a. PCR genotyping confirming the presence of the *GFP-CRF10* transgene.

b. Overexpression of *CRF10* causes an extreme dwarf phenotype. Forty-day-old plants expressing GFP-CRF10 and their segregating wild-type siblings are shown. Scale bar = 1 cm.

## Supplementary Datasets

**Supplementary Dataset 1.** List of eY1H results.

**Supplementary Dataset 2.** Overlap of eY1H and DAP-seq interactions.

**Supplementary Dataset 3.** RSA (root system architecture) phenotyping data.

**Supplementary Dataset 4.** qRT-PCR gene expression data.

## Supplementary Tables

**Supplementary Table 1.** Genotyping, qRT-PCR, and cloning primers.

**Supplementary Table 2.** Summary of mutants, transgenic plants, and promoter-edited plants.

